# Bayesian Optimisation of Large-Scale Biophysical Networks

**DOI:** 10.1101/170779

**Authors:** J. Hadida, S.N. Sotiropoulos, R.G. Abeysuriya, M.W. Woolrich, S. Jbabdi

## Abstract

The relationship between structure and function in the human brain is well established, but not yet well characterised. Large-scale biophysical models allow us to investigate this relationship, by leveraging structural information (*e.g*. derived from diffusion tractography) in order to couple dynamical models of local neuronal activity into networks of interacting regions distributed across the cortex. In practice however, these models are difficult to parametrise, and their simulation is often delicate and computationally expensive. This undermines the experimental aspect of scientific modelling, and stands in the way of comparing different parametrisations, network architectures, or models in general, with confidence. Here, we advocate the use of Bayesian optimisation for assessing the capabilities of biophysical network models, given a set of desired properties (*e.g*. band-specific functional connectivity); and in turn the use of this assessment as a principled basis for incremental modelling and model comparison. We adapt an optimisation method designed to cope with costly, high-dimensional, non-convex problems, and demonstrate its use and effectiveness. We find that this method is able to converge to regions of high functional similarity with real MEG data, with very few samples given the number of parameters, without getting stuck in local extrema, and while building and exploiting a map of uncertainty defined smoothly across the parameter space. We compare the results obtained using different methods of structural connectivity estimation from diffusion tractography, and find that one method leads to better simulations.

## 1. Introduction

Large-scale biophysical models (LSBMs) [40, 35, 4] offer a plausible mechanistic relationship between brain structure (anatomical properties) and function (dynamical properties). This relationship has previously been established by correlating anatomical connectivity (AC) with resting-state functional connectivity (FC) [24, 27, 31], leading to the hypothesis that resting-state activity is an emergent property of the brain, resulting from structured interactions between spatially distributed populations of neurons [16]. As such, it would be one of the few measurable forms of structure-function interaction at the macro-scale, and the ideal activity to compare against large-scale biophysical simulations.

Although the nature of these interactions remains to be characterised, this hypothesis is consistent with more functionally-oriented views, in which the brain is seen as a network of spatially segregated units, co-operating transiently over time in order to carry out the neural computations required for cognition [14, 19]. This view is generally accepted, but still poses many challenges (*e.g.* cortical parcellation, connectome estimation, multimodal integration), some of which affect the large-scale models that we study here. This should be kept in mind when discussing the results obtained with particular models, but the modelling approach itself remains relevant and attractive for many reasons.

Briefly, these reasons pertain either to a methodological, theoretical or clinical perspective. Methodologically, LSBMs offer a unified framework in which previously independent methods – such as diffusion tractography, neuronal population modelling and functional connectivity estimation – are allowed to interact. The ability to connect multiple aspects of brain structure and function via their dedicated fields of study is crucial if we are to build a coherent theory of brain activity. From a theoretical standpoint, these models are designed to provide a mechanistic summary of brain activity in terms of biologically interpretable parameters. A particular model then effectively encodes our understanding of some underlying process, at least to the extent that the empirical data can support. Finally, clinical considerations derive from the theoretical ones; reliable estimates of biologically interpretable parameters can be used to characterise different conditions, or discriminate between them [42].

Here, we focus on the theoretical perspective; specifically with regards to the inference of model parameters from imaging data. Biophysical models typically describe the observed data (*e.g.* fMRI BOLD contrast or MEG) in terms of interpretable parameters (*e.g.* local balance of excitation and inhibition or the hemodynamic response). Because of this formulation, they are *generative* in nature: for a given set of parameters, one can easily generate synthetic data according to the model, which can then be compared to imaging data. However the reverse – estimating the parameters that best fit a given observation, also called *model inversion* – can be very difficult, depending on the number of parameters, the complexity of the model, and the amount of information in the observed data. Unfortunately in practice, empirical estimates of the model parameters are rarely available, and therefore model inversion is required in order to gain insight into the observed data. The main purpose of this paper is to frame inversion of LSBMs as an optimisation problem, propose a powerful method for solving this problem which can handle the computational burden usually associated with simulations, and demonstrate its effectiveness on a simple yet challenging example given the current state-of-the-art.

We model MEG resting-state data using delay networks of oscillatory neuronal masses, with five parameters controlling key structural and functional properties (*e.g.* average delay between brain regions or local frequency responses). This model is formulated mathematically as a large system of non-linear coupled delay-differential equations with over a hundred state-variables, which is numerically delicate and computationally expensive to solve. To further add to the challenges, reliable estimations of functional connectivity patterns (which are compared against empirical measurements from MEG) require on the order of a minute worth of data, and numerical integration methods require timesteps below the millisecond. Therefore, exploring the different ways in which our model behaves as a function of the controlled parameters poses immediate difficulties in terms of computational tractability.

These circumstances call quite naturally for Bayesian optimisation methods [5]; these methods operate under the assumption that the true objective function is computationally expensive to estimate, and instead proceed to *learning* it through iterative cycles of careful exploratory sampling and information consolidation. Specifically, the method presented in this paper is designed for high-dimensional (in practice up to a dozen parameters with typical LSBMs), non-convex and computationally costly problems [30]. It is able to explore the parameter space simultaneously at multiple scales, allowing local optima to compete for the best solution, and using uncertainty estimates to prioritize unexplored regions.

The remainder is organized as follows. First, we present the optimisation method in §2.1 and illustrate the algorithm on a toy-example in Fig. 2. Second, we introduce the LSBM used in our experiments in §2.2, and define the optimisation problem for model inversion (parameters and objective function) in §2.3. The data used in our experiments is described in §3.1, and implementation details are given in §3.2. Finally the results of our experiements are presented in §3.3 and discussed in §3.4.

**Figure 1:**
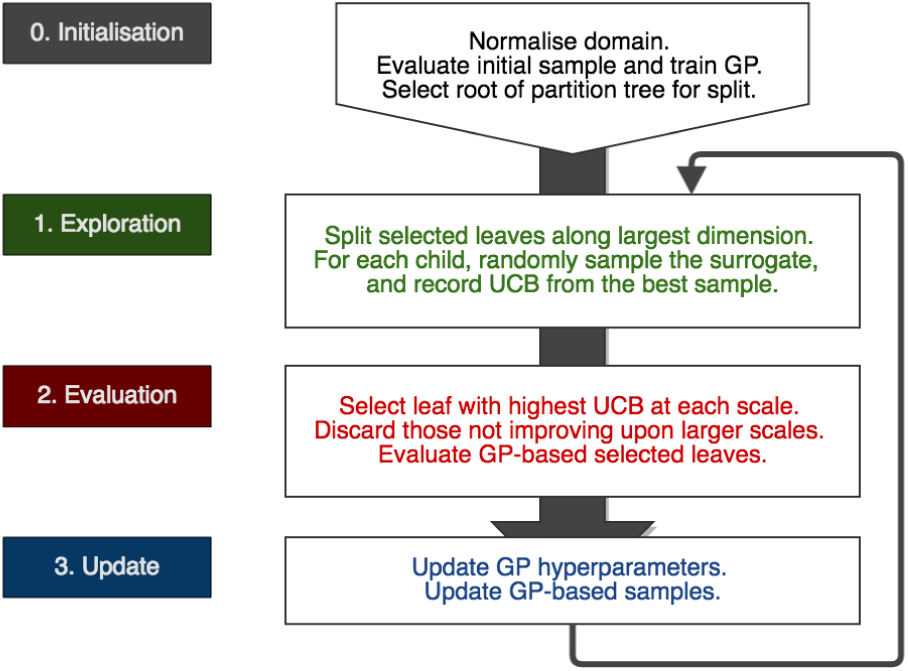
Algorithmic summary of Gaussian-Process Surrogate Optimisation (GPSO). The search space is initially rescaled to normalise the bounds in each dimension to (0, 1). The iterations of the algorithm can be summarised in three main steps; **i)** *exploration*, where selected leaves are partitioned, and children are assessed using GP-UCB; **ii)** *evaluation*, where we evaluate leaves with maximal UCB at each scale, using the objective function; **iii)** *update*, where we re-train the GP including newly evaluated points.

**Figure 2:**
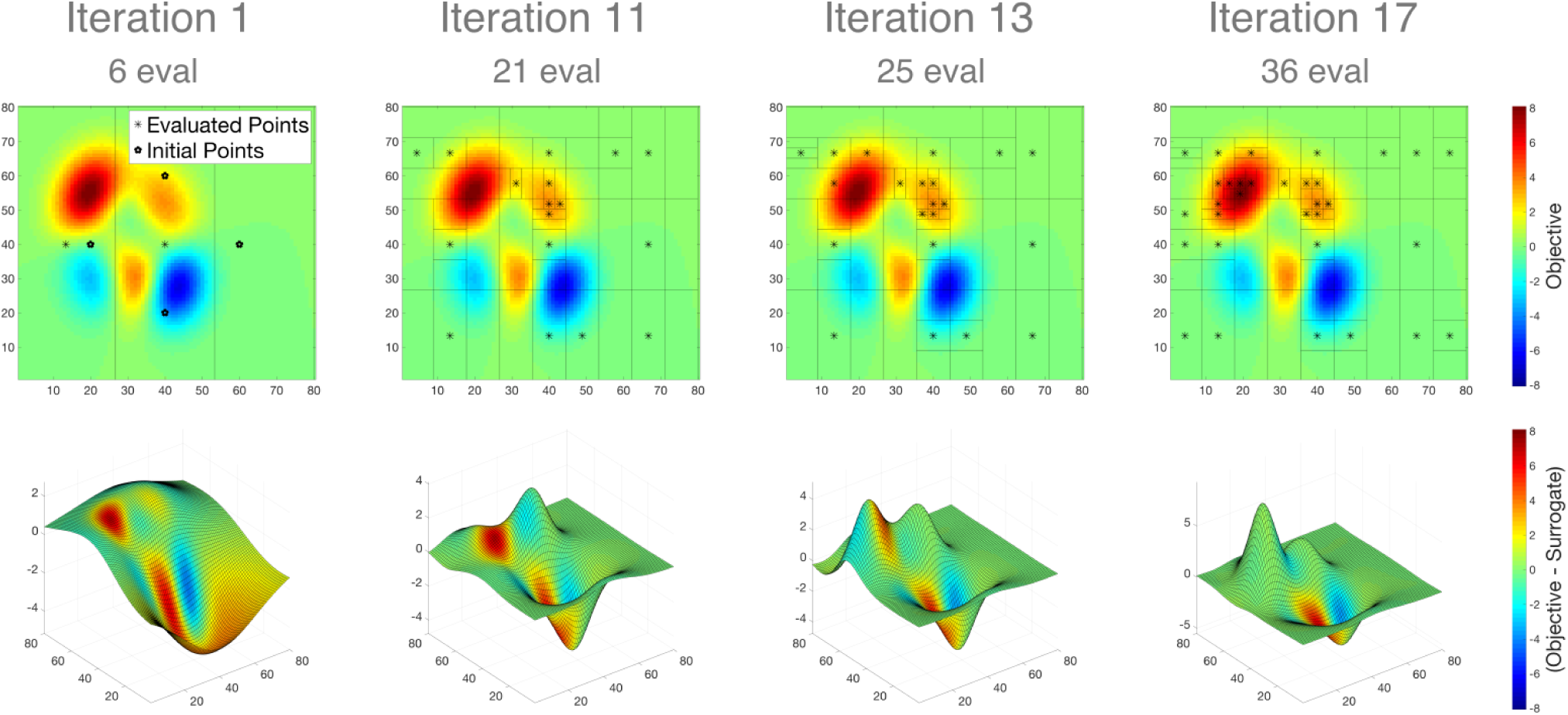
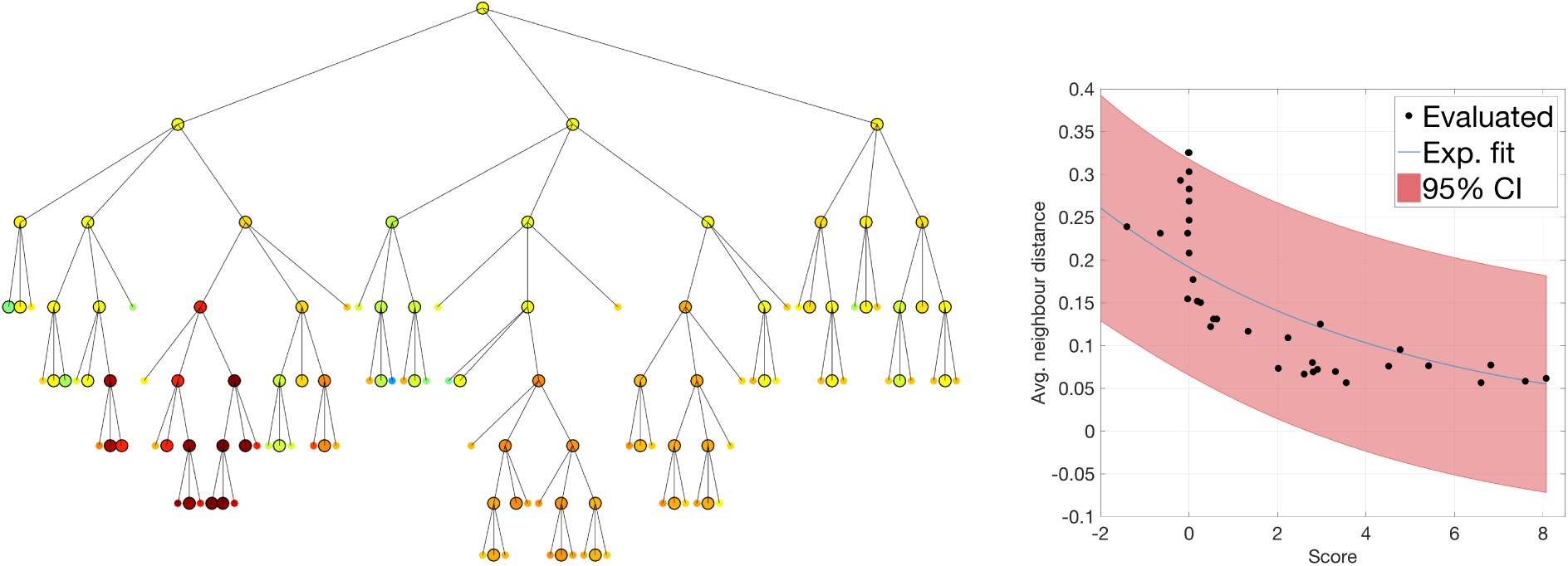
**(a)** Partition (top-row) and surrogate function (bottom-row) for 4 different iterations (columns). Ternary partition (black lines) is shown overlaid on top of the objective function (coloured background). Surfaces show the expected value of the GP surrogate, and colour indicates differences with the true objective: red means true objective > surrogate (conversely for blue). **Iteration 1.** Initial sample and 2 points evaluated in the first iteration; the top and bottom initial points are near a peak and a trough, hence the slope of the surrogate. **Iteration 11.** The algorithm initially finds a local maximum, and converges rapidly to its peak by increasing the number of subdivisions in the area. **Iteration 13.** Exploration at coarse scales hits the slope of the highest peak; surface shows the surrogate peak is misaligned (red patch between the two peaks), but it is already higher than the previous one. **Iteration 17.** Discovery of a higher peak at larger scale froze the subdivision near the first local maximum. The algorithm converged to the global optimum after 4 iterations. The surrogate peak is now aligned with the truth (both peaks are green). **(b) Left.** Ternary partition tree; nodes correspond to subintervals of the search space (see top-row in figure a), colours correspond to the associated scores (upper-confidence bounds), and edges represent set inclusion (parent intervals are the union of their children); in particular, deeper intervals are smaller. Bigger nodes indicate that the objective function was evaluated at their centre, smaller nodes were assessed using GP only. Deeper orange branches at the centre correspond to the local maximum found initially, and red branches on the left correspond to the highest peak. **Right.** Measure of sampling density as a function of the score, showing exponential convergence empirically. For each evaluated point (black dot), the average distance to the 5 nearest neighbours (y-axis, using normalised coordinates) is plotted against the value of the objective function at this point (x-axis). The blue line and red area represent respectively the best fit of an exponential function *x* ↦ *ae^bx^*, and the associated observation bounds with 95% confidence. Gaussian-Process Surrogate Optimisation (GPSO) on Matlab’s `peaks` function.

## 2. Methods

### 2.1. Gaussian-Process Surrogate Optimisation

The method proposed is adapted from [30], and belongs to the family of Bayesian optimisation methods. These methods are designed to tackle computationally expensive black-box global optimisation problems – that is, optimisation problems for which a global solution is sought, but where the objective function is expensive to evaluate, and analytics (*e.g.* the objective’s gradient) are not available. It is worth noting that this method is independent from the particular problem at hand, and may be applied to any other context with similar constraints.

In general, efficient optimisation methods exploit the structural properties of the problem (*e.g.* convexity) in order to devise a strategy which guarantees rapid convergence to a solution. But in the case of blackbox functions, these properties cannot be theoretically determined, and therefore an efficient strategy needs to discover them empirically and adapt as the optimisation progresses. Moreover in the case of expensive objective functions, the strategy needs to restrict the exploration of the search space to a minimum, in order to remain computationally tractable. This excludes in practice all strategies which rely on the gradient or Hessian (because numerical estimates require many function evaluations), but also stochastic sampling methods (*e.g.* MCMC, particle filters or genetic algorithms) which typically rely on large numbers of samples (either for diversity or statistical validity).

#### 2.1.1 Optimism in the face of uncertainty

The problem of finding a suitable strategy given the previous constraints is best formulated within the framework of game theory, where computing-time is seen as a limited resource. The goal is to find the right balance between *exploring* the search space, in order to discover new places of interest with respect to the objective, and *exploiting* the knowledge accumulated by previous iterations, in order to prioritize a more detailed search in places of known interest. This is known as the *exploration*-*exploitation dilemma*, the simplest instance of which is the so-called multi-armed bandit problem (MAB) [2].

In short, the MAB problem consists in picking iteratively from a finite set of possible choices, with repetitions allowed, where the outcome of each choice is random with unknown distribution. For any fixed number of picks, the goal is to maximise the cumulative outcome, by taking the best-known choice as often as possible (exploitation), while regularly trying out unknown or uncertain choices (exploration). A posteriori, the difference between the outcome achieved and the best possible outcome is called the *regret*; minimising the regret or maximising the reward is equivalent.

In this context, a successful balance between exploration and exploitation can be achieved by adopting an *optimistic* strategy, whereby at each turn, the best possible outcome for each choice is considered, given an estimate of uncertainty from previous trials. We then iteratively pick the choice with the best expected outcome, and update our uncertainty according to the result obtained. This strategy is known as the upper confidence-bound method (UCB), and in the next paragraphs we explain how it can be implemented in the context of non-linear optimisation. More detailed explanations about UCB can be found in [7].

#### 2.1.2 Gaussian-Process surrogate

The previous paragraphs give an overview of the strategy adopted, but do not provide a practical solution to our problem. The first issue is that the MAB applies to finite sets of choices, whereas we consider search spaces in which each point is a candidate set of parameters for our models. In fact, adapting the UCB strategy to the latter goes even deeper than considering an uncountable set of choices, it also introduces the notion of a *neighbourhood* for each choice, which should be exploited to enforce smoothness assumptions and propagate knowledge about the objective.

The second issue concerns the representation of this knowledge. Bayesian optimisation methods are only able to tackle such difficult problems because they effectively *learn* the objective as the optimisation progresses, and adapt their search for a solution according to the current state of belief at each iteration. This learned representation is typically defined smoothly across the search space, and much cheaper to evaluate than the true objective function. It can therefore be used as a *surrogate* for the true objective function during optimisation, allowing for computationally tractable analysis and exploration planning. To achieve this, a powerful mathematical tool is required; one not only capable of regressing any sample of points from the objective function (multivariate in general), but also providing smooth estimates of confidence (or uncertainty) across the search space.

Fortunately, this is exactly what Gaussian process regression (GPR) does, and it has been used successfully in the past to solve this second issue [12]. Moreover, resorting to Gaussian processes (GP) also provides intuition into the first issue; GPs can be thought of as an extension of multivariate Gaussian distributions to the infinite case, where any finite subset of points in the search space is itself Gaussian distributed, and the dependence between any pair of points is specified by the *covariance function*, which usually encodes the idea of neighbourhood (typically chosen as a decreasing function of the distance between two points). More details about GPs can be found in [32].

Using GPs, we are able to regress any finite sample of points in order to represent arbitrary objective functions, with an estimate of uncertainty, and with the idea of neighbourhood encoded via the covariance function. The only missing ingredient is a method to overcome the fact that points in the search space cannot be indexed like discrete choices (they are uncountable); without it, the present context of continuous optimisation cannot relate to the MAB problem, and the UCB strategy cannot be applied.

This is achieved in [30] by the introduction of a *partition function*, which splits the search space into distinct subregions that can be explored independently, and can in turn be partitioned themselves to reach a finer resolution – that is, the partition function is *recursive.* Recursivity confers exponential convergence towards regions of interest, and induces a hierarchical structure amongst subregions according to their size (larger regions are non-overlapping unions of the smaller regions contained within them), which can be represented by a *partition tree.* Each node in this tree corresponds to a cartesian region of the search space, covering a unique combination of subintervals in each dimension (*i.e.* a specific range of values for each parameter), and the size of this region decreases strictly with the depth, meaning that we can reach arbitrarily high resolutions. In other words, the partition function allows us to identify regions in the search space with arbitrary resolution, and since there are only a discrete number of nodes at each level, the UCB strategy can be applied in a multi-scale fashion.

#### 2.1.3 Concrete implementation

The main challenge of global optimisation methods, as opposed to local methods, is to deal with local extrema in the objective function. This challenge can be efficiently tackled by carrying out multiple local searches in a sequential (*e.g.* simulated annealing, Metropolis-Hastings) or parallel (*e.g.* particle filters, genetic algorithms) manner. The method proposed here implements a special case of the parallel approach, which organises candidate solutions hierarchically using the partition tree introduced in the last paragraph.

Briefly, the algorithm proceeds iteratively (after initialisation) by: selecting at each level a leaf node with maximal UCB; subdividing selected leaves further using the partition function; exploring children nodes to assess their UCB; and retraining the GP surrogate with new evaluations of the objective function. This is also summarised as a diagram in Fig. 1. Because selected nodes are leaves, we consider at each step a set of regions located in different parts of the search space, and because we select at most one leaf per level in the partition tree, we explore the search space simultaneously at multiple scales.

From there, there are three points to clarify in order to get a concrete implementation:

1. For any point *x* in the search space, the upper-confidence bound is defined as:

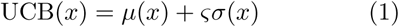

where *μ*(*x*) corresponds to the expected value of the objective function *f* at point *x* given by GPR, *σ*(*x*) is the associated standard deviation, and *ς* is a positive factor controlling our optimism^1^.
2. Each leaf node in the partition tree is labelled as being either: *evaluated*, meaning that the objective function was evaluated at its centre; or *GP*-based, meaning that its associated score was estimated by UCB. Specifically, the score associated with a GP-based leaf corresponds to the best UCB amongst *N* points randomly sampled within the corresponding area in the search space. At each iteration, selected GP-based leaves are evaluated prior to being partitioned, and the score associated with any evaluated node is the value of the objective function at its centre.
3. The partition function is a ternary split along the largest dimension of the subregion considered (in normalised coordinates). This is not a trivial choice; it satisfies several desirable properties with regards to the optimisation, although none of them is required. First, it produces non-overlapping subdivisions, which ensures that there is only one path converging to any specific point in the search space, avoiding redundant competition between nodes. Second, the centre of the parent node is also the centre of the middle child, which saves us an evaluation of the objective function at each split. And third, because of this conserved point, we can guarantee that the children of a node do not recede, meaning that the progression within a branch is monotonic.

Finally, an improvement can be made on the selection process; it is pointless to explore regions at a smaller scale, if some region at a larger scale has a better expected score. Therefore, the selection proceeds sequentially from the root to the deeper branches, and we discard levels at which the maximum UCB does not improve upon the best expected score so far. In effect, this introduces competition between the different scales, and prevents dwelling around local extrema.

### 2.2. Large-Scale Biophysical Model

In this paper, we use the Bayesian optimisation approach introduced in the previous section in order to optimise the parameters of whole-brain dynamical models. Specifically, we consider networks of interacting Wilson-Cowan oscillators with delays. This model posits that the electrophysiological oscillations typically observed in MEG data result from cycles of excitation and inhibition [41], and has been employed previously, notably in [13] to highlight the importance of propagation delays and long-range couplings between distant brain regions, with regards to synchronisation properties in the dynamics produced.

#### 2.2.1 Assumptions and definitions

The brain is modelled as a network of neuronal masses, in which *vertices* correspond to spatially-contiguous brain regions, and *edges* represent direct interactions between these regions. Each neuronal mass may contain several subpopulations of neurons, or several state equations, and so to distinguish between these local entities and the different brain regions in the network, we call *nodes* the vertices corresponding to a subpopulation or state equation, and *units* the groups of vertices located in the same brain region.

We are interested in emergent oscillatory activity in these networks, which is assumed to be driven by cycles of excitation and inhibition in each region. Therefore, two subpopulations of neurons are considered: an *excitatory* subpopulation (E) driving towards increased oscillatory activity, and an *inhibitory* subpopulation (I) driving towards quiescence. The effects of self- and long-range inhibition are neglected, meaning that there are no I-to-I edges, and only E-to-E edges between units. Finally, we do not consider noisy inputs or synaptic plasticity in this paper: their effects has been explored in separate work [1].

#### 2.2.2 Local oscillations

The Wilson-Cowan model [41] describes the temporal variations of the amount of neurons firing within an excitatory and an inhibitory population of neurons, given static local couplings between the two (related to the distribution of synaptic connections), and an external input controlling the excitability of the system.

It introduces so-called “subpopulation response functions”, defined as the cumulative distribution of local firing-thresholds within each subpopulation. These distributions are generally assumed unimodal and symmetric, leading to sigmoidal cumulative functions. In practical terms, the subpopulation response function represents the expected response of an initially quiescent population of neurons to an external input, and is modelled as a logistic sigmoid:

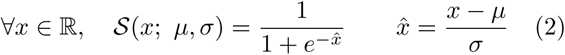

where *μ* represents the response threshold, and *σ* controls the width of the dynamic input range.

Let *E*(*t*) denote the ratio of excitatory neurons firing at time *t* within a brain region (resp. *I*(*t*) for inhibitory neurons). The Wilson-Cowan model states that:

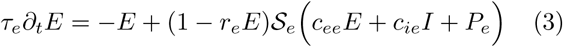

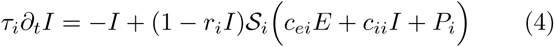

where *∂_t_*• denotes the derivative with respect to time; *c_xy_* = *c*_*x*→*y*_ is the directional coupling of *x* affecting *y*; 𝓢_*e*,*i*_ are the subpopulation response functions; and *P*_*e*, *i*_ are external inputs. The remaining parameters are given in Tab. 1. Notice that although the equations are identical for both subpopulations, the inhibitory coupling coefficients *c_ie_* and *c_ii_* must be non-positive (by definition), while the excitatory coefficients *c_ee_* and *c_ei_* must be positive, which breaks the apparent symmetry between excitation and inhibition.

Applying our assumption about inhibitory self-coupling, we set *c_ii_* = 0. Furthermore, given that the refractory periods are typically much smaller (~ 10^−3^) than the scale of variation of the state variables (interval [0, 1]), their effect in practice is negligible at such large scales and therefore we set *r_e_* = *r_i_* = 0. In summary, the oscillatory mechanism of this model is simple: i) excitatory inputs lead to an increase in excitatory activity; ii) excitatory activity causes an inhibitory response; iii) decreased excitation leads to a decreased inhibition; iv) decreased inhibition leads to a relative increase of excitatory inputs.

The architecture of this model, as well as the typical dynamics produced, and the effect of key local parameters on these dynamics, are shown in Fig. 3.

**Figure 3:**
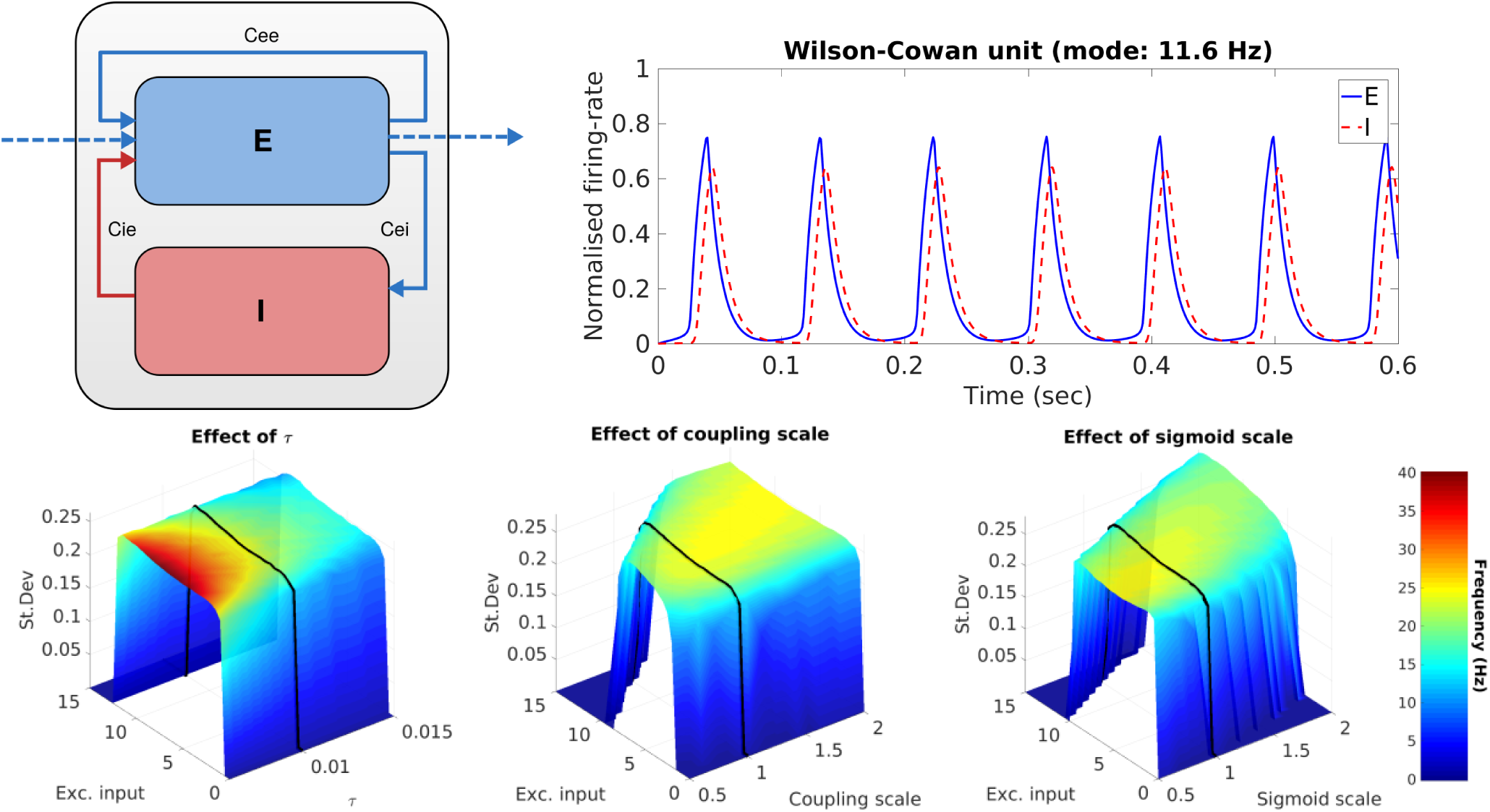
Illustrations of the Wilson-Cowan unit. **Top-left:** local two-population structure (excitatory and inhibitory), without self-inhibition (*c_ii_* = 0) and with long-range excitation only (blue dashed lines). **Top-right:** example oscillatory timecourse showing inhibition (red dashed line) lagging behind excitation (blue plain line); the lag is controlled by the time-constants *τ*_*e*,*i*_, and here the excitatory input is set to 0.84. **Bottom-row:** evolution of standard-deviation (surface height) and frequency mode (colormap) as a function of the excitatory input, and varying parameters in three different ways. Black lines correspond to an increasing *P_e_* with the baseline parameters (see Tab. 1). The unit is always silent without excitatory input, and saturates for large inputs – the interval between oscillatory and saturation thresholds is the *dynamic range* of the unit. Notice that the frequency of oscillations depends on the input; this property allows remote brain regions to affect the local phase via their connection, which is a potential mechanism for long-range synchronisation. *Left:* the frequency of oscillations can be controlled with the time-constant *τ* without affecting the dynamics. *Middle:* an upscale of local couplings dilates proportionally the dynamic range of the unit. *Right:* small scalings of the response parameters (*μ*, *σ*) linearly translate the dynamic range, but also affect the range of oscillatory frequencies.

**Table 1:** Baseline parameters for the Wilson-Cowan model (see Eq. 3,4). Where subscripts are omitted, the description and value of the parameter apply to both subpopulations. The excitatory input *P_e_* is controlled during our experiments. The response parameters (*μ*, *σ*) were set such that small inputs (compared to the dynamic range, see Fig. 3) would cause the system to oscillate. Couplings were set according to a ratio of 80% self-excitation (*c_ee_*/(*c_ee_* + *c_ei_*) = 0.8), and no self-inhibition (*c_ii_* = 0).

| Symbol | Description | Value |
| --- | --- | --- |
| $\tau$ | Time-constant | 10 ms |
| $r$ | Refractory-period | 0 ms |
| $\mu$ | Response threshold | 3 |
| $\sigma$ | Dynamic range | $\mu/6$ |
| $c_{ee}, c_{ei}$ | Excitatory coupling | 28, 7 |
| $c_{ie}, c_{ii}$ | Inhibitory coupling | -35, 0 |
| $P_i$ | Inhibitory input | -0.3 |

#### 2.2.3 Network extension

Extending the previous local equations to a network of interacting brain regions consists in adding coupling terms from those remote regions inside the subpopulation response functions. The general *node* equation (whether excitatory or inhibitory) in a network of *N* brain *units* is therefore:

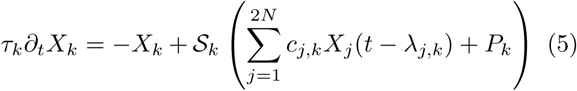

where 1 ≤ *k* ≤ 2*N* with the convention that odd indices correspond to excitatory nodes (resp. even for inhibitory nodes); *X_k_* is the normalised firing-rate of node *k* (corresponding to previous variables *E* and *I* at the unit-level); and we introduced delay parameters λ_*j*,*k*_ ≡ λ_*j*→*k*_ ∈ ℝ_+_ to account for propagation times between distant brain regions. These delays are of the same order of magnitude as the characteristic time-constants of local subpopulations, and therefore interfere with their dynamics^2^.

### 2.3. Model Optimisation

The model presented in the previous section describes the activity of a network of *N* brain regions, using 2*N* state equations (see Eq. 5). In general, this network will not be sparse, meaning that there are 𝓞(*N*) non-zero coupling terms in most state equations, hence the high computational costs associated with simulations in practice (there are 𝓞(*N*^2^) interaction terms to be computed at each time-step). As it stands, there are also 𝓞(*N*^2^) parameters, because of the coupling and delay matrices, respectively [*c*_*i*,*j*_] and [λ_*i*,*j*_]. It is therefore impractical to move on directly to the simulation of such systems, without a more parsimonious parametrisation of the model.

In this section, we propose a simple parametrisation controlling key structural and functional aspects of the system with few parameters. These parameters can be inferred from empirical MEG data, using the method presented previously in §2.1, by framing model inversion as an optimisation problem, for which we propose an objective function below.

#### 2.3.1 Assumptions

For simplicity, we assume that all units in the network are identical, and that excitatory and inhibitory sub-population response functions and time-constants are identical (see Tab. 1 for baseline parameters). Each unit is normally defined by 9 parameters (*τ*, *μ*, *σ* for each node, and *c_ee_*, *c_ei_*, *c_ie_*), so these assumptions reduce the number of unit parameters from 9*N* to 6.

Since there are two nodes per unit (excitatory and inhibitory subpopulations), the connectivity and delay matrices have a 2-block structure. For instance, with the coupling matrix, all on-diagonal blocks are identical (and contain the local couplings), and off-diagonal blocks only have one non-zero entry (only E-E long-range connections):

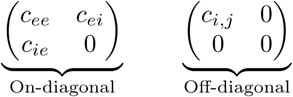

With the delay matrix, we reason in pairs of units instead of nodes (*i.e.* the delay between two regions is the same regardless of which subpopulations we consider in each). Therefore the 2-block between units *i* and *j* is simply:

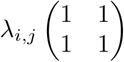

and we neglect delays within units (λ_*i*,*i*_ = 0). Delays are estimated from pairwise Euclidean distances, and we assume a constant propagation velocity throughout the brain to avoid introducing additional parameters.

Furthermore, we only consider cortico-cortical connections in this work, and assume that the two hemispheres correspond to subnetworks of equal size (*N*/2 units). The latter induces an additional N-block structure in the previous matrices, which is useful for two reasons:

- to our knowledge, there is no evidence for one hemisphere driving brain activity more than the other, or for a lateral bias in the AC between hemispheres, therefore requiring both to have the same size ensures that the overall AC within and between hemispheres is structurally unbiased;
- from a purely practical perspective, the assumption of hemispheric symmetry makes it easier to manipulate connections within and between them, as in Eq. 8 for instance.

Finally, note that despite these numerous assumptions the network is still heterogeneous due to the different coupling weights and delays assigned to the edges of the network; this is consistent with the overall objective of studying the effects of structural properties on dynamical activity.

#### 2.3.2 Parametrisation

Let *D* be the matrix of pairwise Euclidean distances between brain regions, and *A* the associated matrix of anatomical connectivity estimated from diffusion tractography (both *N* × *N*). By convention, the diagonal of *A* is set to zero, and we recall that excitatory and inhibitory nodes are indexed between 1 and 2*N*, respectively with even and odd numbers.

The coupling matrix *C* = [*c*_*i*,*j*_] and delay matrix Λ = [λ_*i*,*j*_] are parametrised respectively as follows:

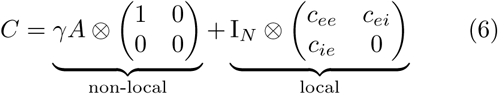

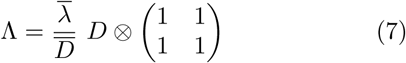

where ⊗ is the Kronecker product; I the identity matrix; *D̅* the average pairwise distance; and we introduced the following parameters:

- *γ* the global coupling strength, controlling the overall amount of non-local coupling;
- and λ̅ the average propagation delay, controlling the speed of interactions.

Note that although matrix *A* might be symmetric, *C* is *not*; the element in row *i* column *j* corresponds to the edge from node *i* to node *j* (not unit), and therefore each column can be seen as a coupling vector for the corresponding node.

Probabilistic tractography methods have an inherent bias towards shorter connections; longer streamlines are less probable, and therefore connectivity between distant regions is generally lower [37] (see Fig. 4). This reflects a biological reality [18], but beyond the issue of assessing the accuracy of the estimated decrease, there is the question of whether the same decrease rates apply equally within or between hemispheres. In order to correct for such potential bias, we introduce an additional parameter *h* to manually scale inter-hemispheric connections, which correspond to the off-diagonal N-blocks in matrix *C*. This scaling is affected to *A* directly, before substitution in Eq. 6:

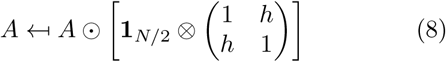

where ⊙ is the Hadamard product (element-wise) and **1** is a full matrix of ones.

**Figure 4:**
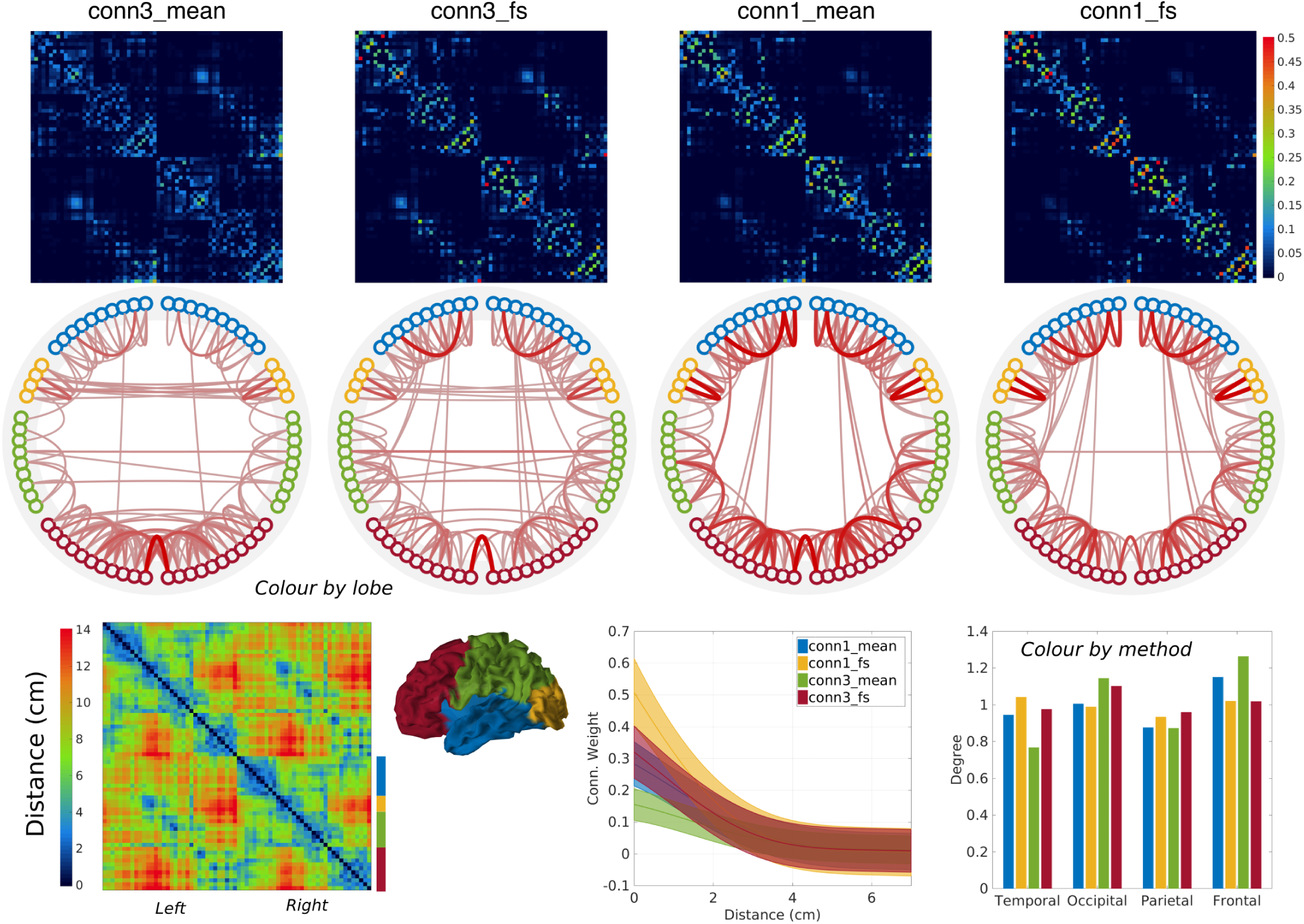
Structural information used in the biophysical models. **Row 1:** AC matrices estimated from diffusion tractography, using two different seeding methods (conn1, conn3), and two normalisation methods (mean, fractional scaling), see §3.1.1 for details. **Row 2:** thresholded network (90^th^ percentile) showing the strongest edges in corresponding AC matrices. conn3 seeding favours homotopic connections, whereas conn1 favours anterior-posterior connections, and mean normalisation shows stronger connectivity in the frontal lobe. **Bottom-left:** matrix of pairwise distances showing hemispheric block structure. Lower distances around the diagonal are due to the ordering of the different regions (chosen manually). **Bottom-right:** basic statistics on connectivity weights. Connectivity decreases exponentially with the distance (left, GP regression showing predicted means and 95% confidence intervals). Average degrees are higher in the frontal and occipital lobes (right, bars shown for each method, and grouped by lobe); fractional scaling reduces frontal connectivity, while increasing temporal and and parietal ones; and `conn1` seeding yields noticeably higher connectivity in the temporal lobe, and lower in the occipital lobe.

Finally, we consider two functional parameters affecting the oscillatory dynamics of all units:

- the time-constant *τ*, assumed equal for all nodes, which controls the frequency response of Wilson-Cowan units (see Fig. 3);
- and the excitatory input *P_e_*, assigned equally to all excitatory nodes across the network, which controls the *excitability* of individual units when they are below oscillatory threshold.

Equations 6, 7 and 8 determine entirely the network structure, and we consider *five parameters* to be optimised, in Tab. 2, which control key structural and functional aspects of our model.

**Table 2:** Network parameters controlled during optimisation. The ranges correspond to the boundaries of the search space (required by GPSO). The parameter variants *P͠_e_* and *γ͠* are defined in §2.3.3. Short names are used in figures 6, 9 and 11.

| Symbol | Description | Short-name | Range |
| --- | --- | --- | --- |
| $\widetilde{P}_e$ | Relative input | Input | (0.6, 1) |
| $\widetilde{\gamma}$ | Relative coupling | Coupling | (1, 3) |
| $\bar{\lambda}$ | Average delay (ms) | Delay | (1, 50) |
| $h$ | Inter-hem. scaling | IH Scaling | (0, 4) |
| $\tau$ | Time-constant (ms) | Tau | (4, 16) |

#### 2.3.3 Relative variants

The previous parameters control key structural and functional aspects of our LSBM, but their range of values can vary depending on the AC matrix considered (and more generally, the oscillatory unit considered). This means that a suitable domain for optimisation needs to be determined *ad hoc* every time, which makes it difficult to compare solutions found across models.

We know (see Fig. 3) that Wilson-Cowan units oscillate for excitatory inputs beyond a certain threshold value 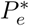. Similarly at the network level, we know that oscillations occur for coupling values beyond a certain threshold value *γ*^∗^ (which is null if the units intrinsically oscillate on their own).

Normalising these parameters with respect to their threshold value would help, not only to compare them across different models, but also to easily control the state of the network (oscillating or silent) and focus on the oscillating regimes during optimisation. Hence, we define the following *relative* variants instead:

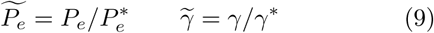

and use them throughout our experiments.

With these definitions, we know for example that *P͠_e_* < 1 corresponds to brain units below oscillatory threshold, and that networks are in oscillatory regime only when *γ͠* > 1. And we can enforce these conditions during optimisation by choosing the parameter ranges accordingly (see Tab. 2).

However, determining the threshold value *γ*^∗^ is not trivial, because it depends on 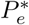 (the unit oscillatory threshold), as well as on other controlled parameters such as the average delay and inter-hemispheric scaling. While 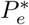 can be determined numerically (*e.g.* with bifurcation analysis), to our knowledge there is no simple method for estimating the oscillatory coupling threshold *γ*^∗^ for any given delay-network.

In our experiments, for any candidate set of parameters (including normalised input and coupling), both threshold values were estimated prior to simulation in order to determine the corresponding values *P_e_* and *γ*, which are required in order to build the network (see previous section). This was done by dichotomic search with a precision of 3 significant digits. The overhead introduced, in terms of runtime, was on the order of a minute per candidate set of parameters (largely dominated by the search for *γ*^∗^; the search for 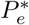 always took less than a second).

#### 2.3.4 Objective function

The *optimal* parameters should maximise the similarity between biophysical simulations and real MEG data, and this similarity should be assessed using characteristic features of resting-state dynamics. In this paper, we take a simple objective function comparing FC matrices across six overlapping frequency bands:

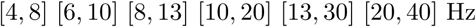

As excitatory pyramidal cells contribute most strongly to EEG/MEG signals, we associate activity in the excitatory populations of the model with signals in experimental data [8]. Envelope correlations were computed in each band, as is commonly done with resting-state MEG (more details in §3.1.2). Importantly, the simulated timeseries were orthogonalised prior to computing Hilbert envelopes (using the Procrustes method from [11]), in order to replicate the effects of leakage correction on source-reconstructed MEG data.

Denoting *M*_1.6_ the corresponding FC matrices, where subscripts identify the frequency-band, we define the vector of *relative connectivity magnitudes* as:

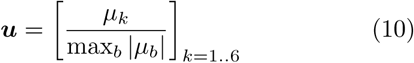

where *μ_b_* is the average off-diagonal correlation coefficient in matrix *M_b_*. By definition, the largest element in this vector has magnitude 1 (*e.g.* in alpha band), and the magnitude of each element gives the amount of connectivity in one band compared to the principal one (*e.g.* in theta compared to alpha).

This vector is computed for the simulated and reference data independently, in order to compare the relative amounts of connectivity across frequency bands.

Note that because we divide by the largest correlation coefficient across bands, this comparison is insensitive to any scaling of either set of matrices (reference or simulated), which can vary as a function of the signal-to-noise ratio for instance, or the amplitude of the oscillations.

Finally, the objective function used in our experiments combines the similarity between relative connectivity magnitudes, and the average within-band correlation between simulated and reference FC matrices:

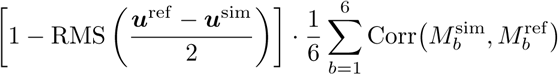

where superscripts refer to the simulated or reference data, and the first factor is a normalised measure of similarity (in [0, 1]) based on a root-mean-square metric, which is 1 when *u*^ref^ = *u*^sim^, and decreases towards 0 as the distance between them increases.

## 3. Results & Discussion

### 3.1. Imaging data

#### 3.1.1 Anatomical structure

The Desikan-Killiany cortical parcellation [17] was used in all experiments to define brain regions (or “units” in our network models). The AC between regions was estimated using probabilistic diffusion tractography [3, 28], and averaged across 10 diffusion MRI datasets from the Human Connectome Project (HCP) [39, 36]. Distortion corrected data [22] was used to estimate fibre orientations [29, 26], and used subsequently for probabilistic tractography in FSL. Delays between regions were estimated using Euclidean distances between the region’s barycentres.

Two different seeding methods were used to compute dense tractography connectomes: with the conn1 method, streamlines were seeded from the WM/GM interface; whereas the `conn3` method considered every brain voxel as a seed. The number of streamlines reaching locations on the WM/GM boundary (~60k vertices in standard MNI space, as given by the CIFTI format [22]) were recorded.

Both connectomes were then parcellated and normalised in order to estimate anatomical connectivity between each region. Two different normalisation methods were used [18]:

- the `mean` method counts the number of streamlines between pairs of vertices belonging to two regions, and divides by the number of vertices in both;
- whereas fractional scaling (`fs`) divides instead by the sum of the source-region output count and target-region input count.

Conceptually, the first normalisation accounts for differences in *size* between different regions, while the second method accounts for differences in *connectivity* between pairs of regions instead (which indirectly accounts for differences in size as well).

Finally, each connectivity matrix was made symmetric by arithmetic average with its transpose, and rescaled such that the average degree (sum of rows or columns) be unitary. The corresponding AC matrices are shown in Fig. 4.

#### 3.1.2 MEG resting-state

The resting-state datasets of 28 healthy subjects from [6, 33] was used in our experiments. Details about the acquisition and pre-processing can be found in these references. The data were beamformed into MNI 8mm standard space between 4 and 40Hz, parcellated using PCA, rescaled to set the largest standard-deviation to 1, and orthogonalised to correct for spatial leakage using the Procrustes method from [11].

Each dataset was then filtered in the following six overlapping frequency bands:

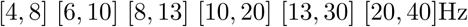

and correlations between Hilbert envelopes were computed in each band. The resulting band-specific FC matrices were then averaged across 28 subjects, and taken as *reference data* for our simulations to be compared against. These reference FC matrices in theta, alpha and beta bands are shown along with the best simulated results in Fig. 8.

In addition, we performed a time-windowed analysis on real MEG data in order to assess the best similarity scores to be expected as a function of the simulation time-span in our experiments (see objective function in §2.3.4). Specifically, for a window of a given time-length, we extracted segments of source-reconstructed time-series from all 28 MEG datasets, estimated the functional connectivity matrices for each of these segments, and computed the associated similarity scores as if those were simulated data. The distribution of scores obtained (see Fig. 5) was taken as a gold-standard for our simulations; we should expect our best simulations to hit the upper-end of this distribution, but significantly higher scores would indicate overfitting, and lower scores would indicate poor model performance.

**Figure 5:**
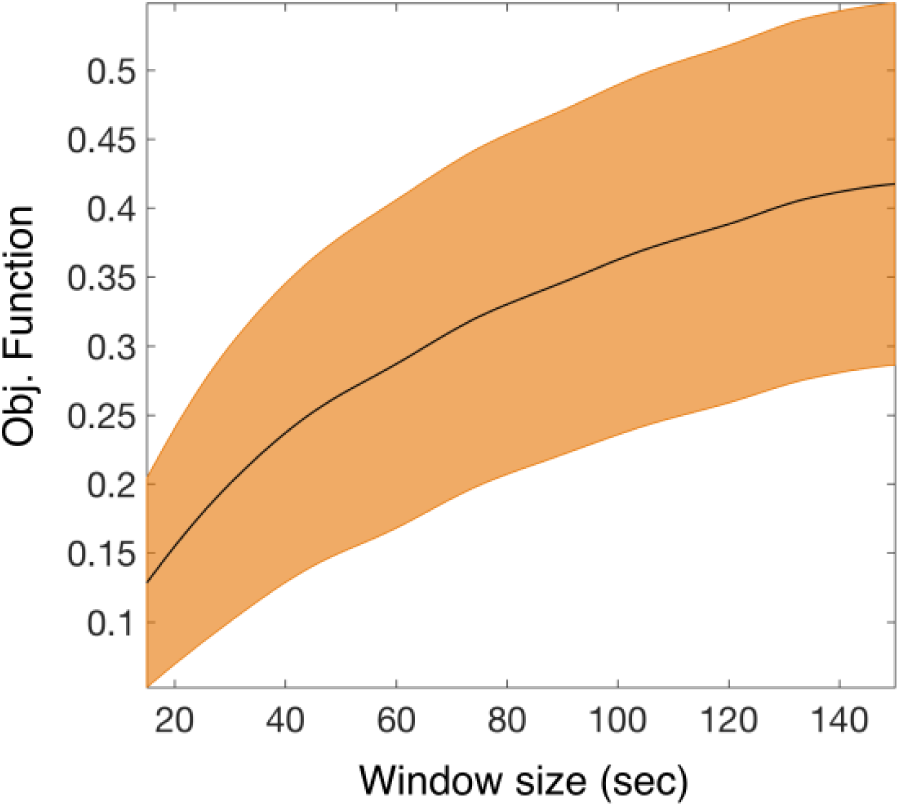
Distribution of similarity score (see §2.3.4) estimated on real MEG data across all 28 subjects, for time-windows of varying length (15 to 150 sec, 50% overlap). The black centreline is the median similarity score as a function of the window-length, and the orange patch shows the associated 95% confidence interval. This distribution is used as a gold-standard to assess the performance of our simulations; for a time-length of 60 sec, the upper similarity bound with 95% confidence is 0.41, and the best score obtained with our simulations is 0.42 (see Fig. 10).

We also used this analysis in order to strike a reasonable balance between higher expected scores and longer simulation times. The computational costs associated with longer simulations were considerable, and this analysis allowed us to assess the expected penalty for choosing shorter simulation times. We opted for simulations with an equivalent of 60 seconds worth of data in our experiments (downsampled to 300 Hz before analysis); for this time-length, the corresponding upper-bound for the expected similarity scores with 95% confidence is 0.41, and the best score obtained in our experiments was 0.42 (see Fig. 10).

### 3.2. Software implementation

#### 3.2.1 GP Surrogate Optimisation (GPSO)

We improved upon the implementation of IMGPO [30], by addressing a number of issues and extending the algorithm in several ways. Our implementation is a complete refactoring of the original algorithm, and is made freely available under the terms of license AGPLv3^3^ at the following address^4^: https://gitlab.com/jhadida/gpso.

Our main contributions are listed below:

- to update upper-confidence bounds following the optimisation of GP hyperparameters at each iteration, in order to allow belief propagation across the partition tree;
- to enable the exploration of candidate leaves using uniformly random samples of points in the corresponding subregion of the search-space (the original implementation only explored a subset of the dimensions in a deterministic manner);
- to implement serialisation, allowing for the optimisation to be resumed at any stage.

The various settings used during our experiments are listed in Tab. 3.

**Table 3:** GPSO hyperparameters and initial values used for all experiments. The optimism parameter *ς* corresponds to confidence bounds of 99.5% (*i.e.* erfc^−1^(0.005)), which was found to strike a good balance between exploration and exploitation.

| Function | Hyperparameter | Value |
| --- | --- | --- |
| UCB | $\varsigma$ | 1.98 |
| Constant mean | $\mu$ | 0 |
| Gaussian likelihood | $\sigma$ | 0.001 |
| Isotropic Matérn covariance (order 5) | Length | 0.25 |
|  | Magnitude | 1 |

#### 3.2.2 Biophysical Simulations

The LSBM presented in §2.2 was implemented in C++, and simulations were analysed with Matlab.

The system of non-linear coupled delay-differential equations (see Eq. 5) was solved using an adaptive-step Runge-Kutta method of order 8 adapted from the reference Fortran implementation `Dopr853` in [25]. The main computational bottleneck in the simulations is due to the number of feedback terms to be computed at each time-step; since network matrices (delay and coupling) are not sparse, the complexity is quadratic in the number of nodes in the network. At each timestep of size *h*, the sum of delayed terms in each equation were computed across multiple threads at time *t* and *t* + *h*, and interpolated for each substep using an exact formula (that is, the interpolation does not make any approximation). These optimisations allowed for simulation times roughly two times slower than real-time using four threads on modern CPUs.

The initialisation of delay-systems is delicate. In contrast with initial value problems, which typically require a single initial state, delay-systems require a smooth *function* for initialisation. This function must be defined over a time-interval [*t*_0_ – λ, *t*_0_], where *t*_0_ is the initial time and λ is at least as large as the largest delay. Additionally, it should itself be a solution of the system, which makes the problem circular.

To our knowledge, there is no solution to this problem. In our experiments, for each simulation, we calculated the fixed point (*E*, *I*) to which individual units converged given the current excitatory input^5^, and set the initial function to be constant and equal to these values in each unit. It is equivalent to assume that units are initially disconnected from the network for a certain period of time.

### 3.3. Experiments

We present the results of two experiments which demonstrate the benefits of GPSO in the context of LSBMs. The first experiment is a proof of concept in a restricted two-dimensional case, which allows results to be visualised and compared with exhaustive search. The second experiment considers the full model with five parameters, for which we provide a detailed analysis of the results and highlight the current limitations.

#### 3.3.1 Two-dimensional example

In this experiment, the similarity between simulated and reference MEG data was maximised according to the objective function defined in §2.3.4, by optimising just two parameters for now; the average delay λ̅, and the relative network coupling *γ͠*. The remaining parameters (see Tab. 2) were set to: *P͠_e_* = 0.85, *h* = 1, *τ* = 10ms, and we used the `conn1_mean` AC matrix to connect the network units.

The timespan of each simulation was 63 seconds, and we discarded the first 3 seconds to get rid of transient effects before analysis. The results are shown in Fig. 6.

**Figure 6:**
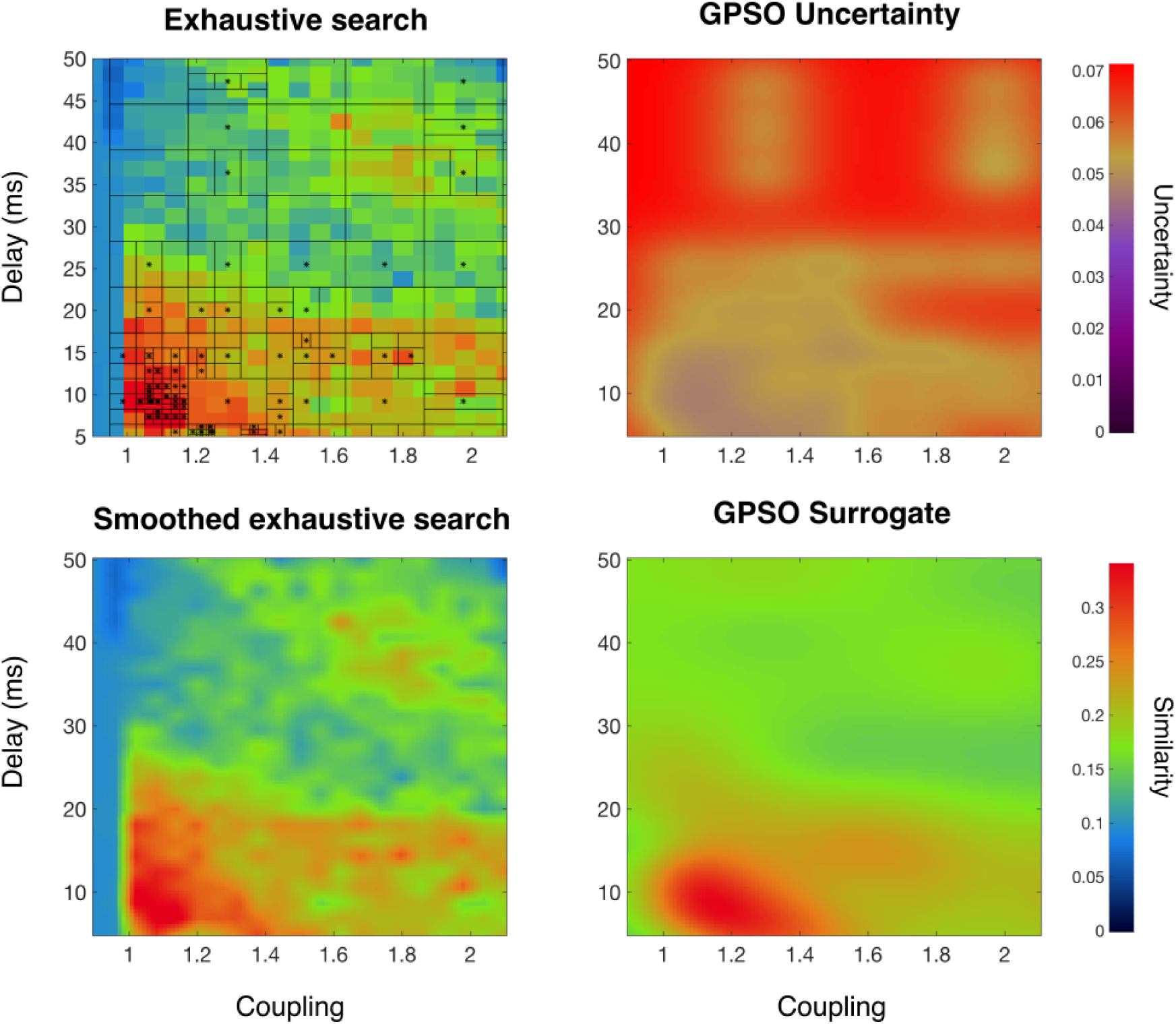
Exhaustive grid-search with 525 simulations, compared against GPSO with 100 simulations, controlling 2 parameters (average delay and relative coupling). The remaining parameters (see Tab. 2) were set to: *P͠_e_* = 0.85, *h* = 1, *τ* = 10ms, and we used the `conn1_mean` AC matrix to connect the network units. **Top-left:** exhaustive search (background image) and partition tree from the GPSO (black lines). Black asterisks indicate the samples evaluated during optimisation (see §2.1.3 for details about GP-based samples). Each pixel corresponds to a 63 sec simulation, analysed and compared with reference MEG data. The partition is refined in places where the objective function is higher, and the optimisation converged rapidly to the global optimum. **Bottom-row:** surrogate function (predicted mean) learned by GPSO, to be compared against the smoothed exhaustive search (ground-truth) on the left. **Top-right:** surrogate uncertainty (predicted st-dev.), driving the compromise between exploration and exploitation during optimisation.

The performance of GPSO was assessed by comparison with an exhaustive grid search, which is computationally tractable with two dimensions and can be easily visualised. The grid search required 525 simulations, considering respectively 25 and 21 equally spaced points across the value ranges of the delay and coupling parameters. In comparison, GPSO was run with 100 simulations, with which it successfully converged to the optimum, while learning a surrogate objective function defined smoothly across the search space, along with a map of uncertainty. These results demonstrate the efficiency of the method in a restricted two-dimensional context of LSBM optimisation.

#### 3.3.2 Five-dimensional analysis

In this second experiment, we consider all five parameters listed in Tab. 2, and all four connectivity matrices shown in Fig. 4. For each connectivity, an optimisation was run with 800 samples (*i.e.* evaluations of the objective functions), which took approximately 1.5 day to run on a computing cluster with four threads. In comparison, an exhaustive search run sequentially with just 20 values per dimension would take over 18 years to complete.

The five-dimensional results cannot be displayed as in the previous two-dimensional case; instead we summarise below key aspects of the analysis, illustrating the type of information made available by this new method.

**A case for multi-criteria objective functions** • Defining the “goodness-of-fit” with resting-state electrophysiological data is a difficult task, especially given the time-constraints typically associated with LSBM optimisation. Here, we discuss the benefits of including a penalty factor in the objective function, to ensure that the relative amounts of FC across frequency bands are similar in real and simulated data. It is best to have the main points of §2.3.4 in mind when reading this paragraph.

We illustrate our point in Fig. 7, where the best results obtained after optimisation with each of the four AC matrices are summarised and compared. Without the penalty term included in the objective function to control for the relative strength of connectivity across frequency bands, the results obtained with `conn3_fs` connectivity were better than those with `conn1_mean` connectivity, despite the fact that the corresponding FC matrices (see bottom-row in Fig. 8) are almost identical across frequency bands, and only vary slightly in terms of connectivity scale.

**Figure 7:**
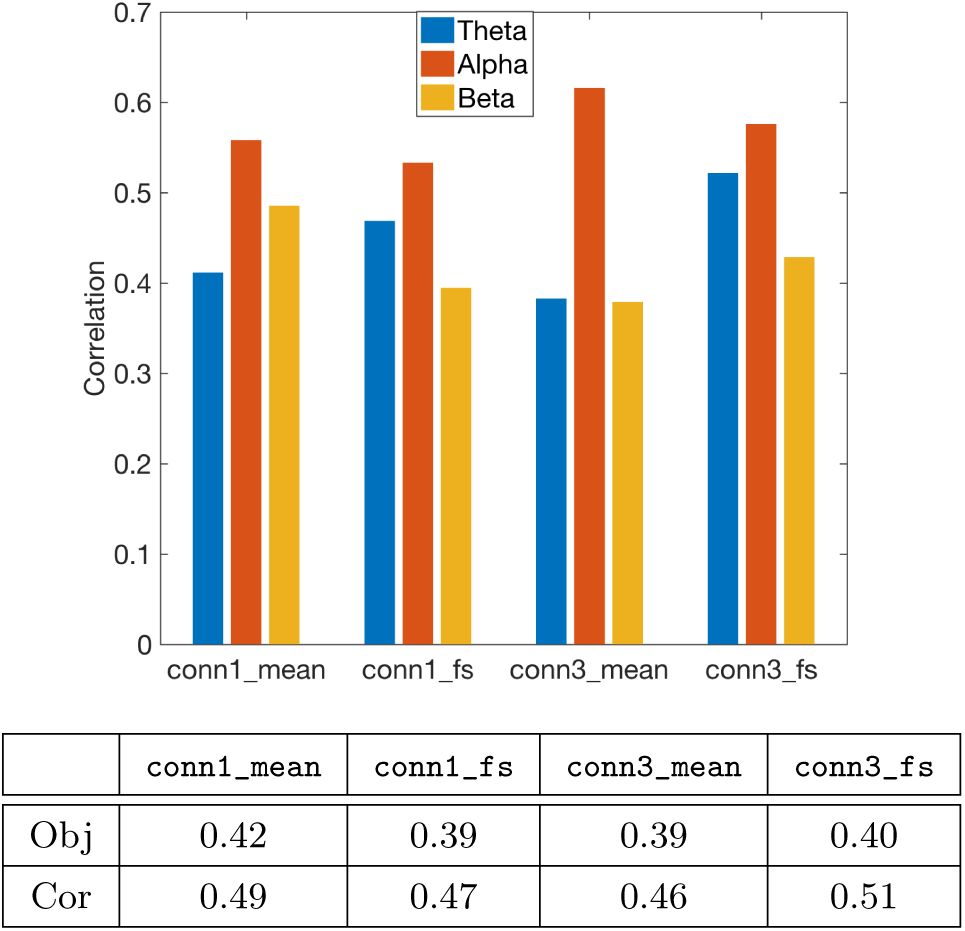
Comparison of best results obtained with the four AC matrices shown in Fig. 4. The bar plot shows the correlation between simulated and reference FC matrices in theta, alpha and beta bands. The table reports the average correlation in these bands, as well as the similarity score calculated with the objective function in §2.3.4. Without the penalty term included in the objective function to control for the relative strength of connectivity across frequency bands, the results obtained with `conn3_fs` connectivity were better than those with `conn1_mean` connectivity, despite the fact that the corresponding FC matrices (see bottom-row in Fig. 8) are roughly identical across frequency bands. This illustrates the importance of choosing a suitable objective function.

**Figure 8:**
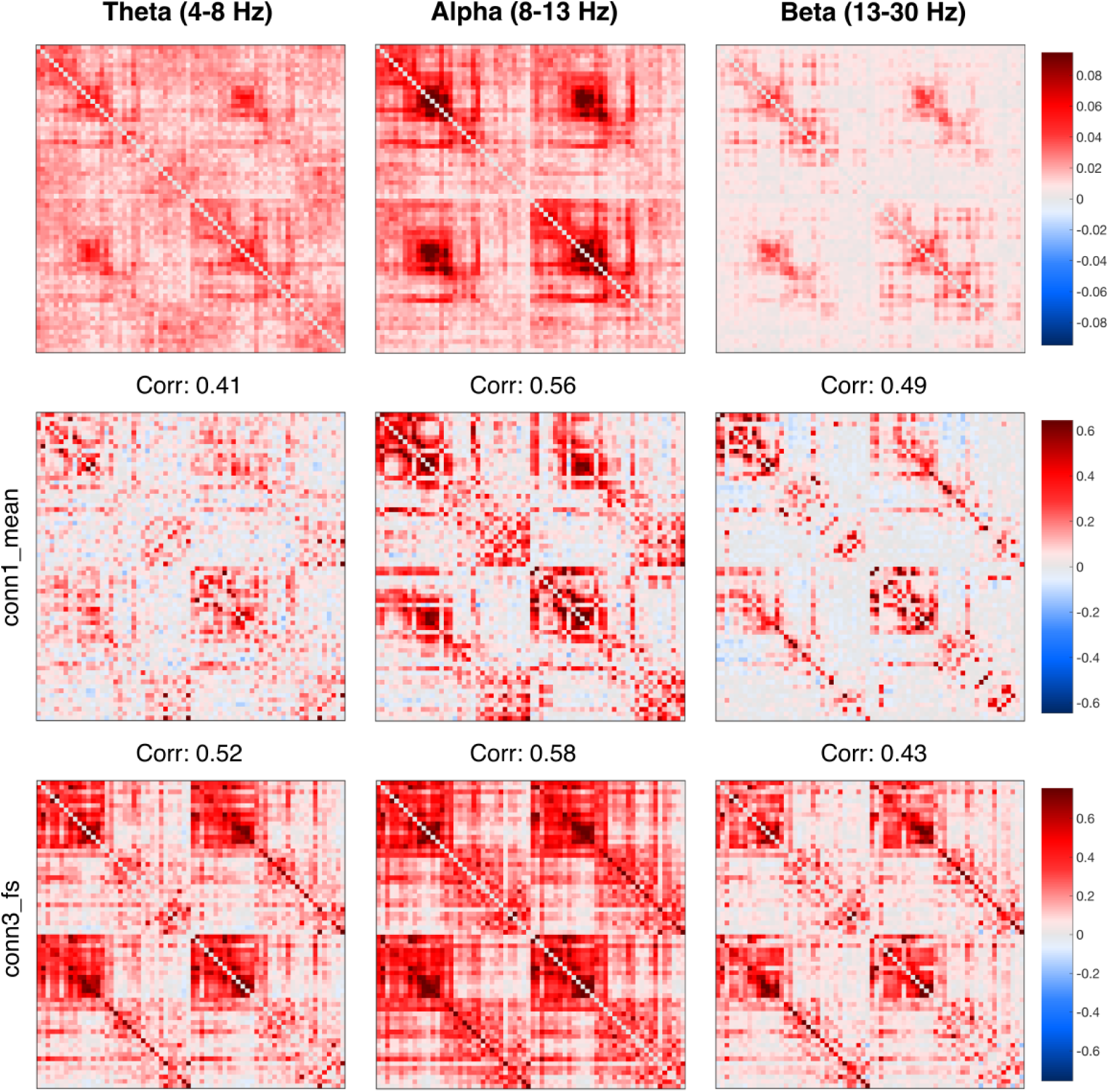
Comparison between simulated and reference FC matrices in theta, alpha and beta bands. Reference matrices are shown in the first row, followed by the best results obtained with connectivity `conn1_mean` (row 2), and the second best results obtained with `conn3_fs` (row 3). The correlation between each simulated FC matrix and the corresponding reference is indicated on top of the matrix. The FC patterns obtained with `conn1_mean` connectivity are strikingly similar to the reference, except in the frontal lobe (lower-right block in each quadrant). Note that although results obtained with `conn3_fs` achieved better correlations on average, they had a lower similarity score than the results obtained with `conn1_mean`, because their variation across bands was poor (see Fig. 7).

The FC matrices obtained with `conn1_mean` connectivity also had a better structural correspondence with the reference matrices (see top rows in Fig. 8), but this was only by chance; the penalty term did not favour this correspondence in any way. In fact, this is one of the weaknesses of the correlation coefficient itself, which does not take into account structural dependencies between the elements of the FC matrices (*i.e.* the connectivity *patterns*) when comparing them.

To summarise, these results demonstrate that the inclusion of a penalty term controlling for relative strengths of FC across frequency bands was beneficial in our experiments, and suggest that multi-criteria objective function might in general be desirable in the context of LSBMs. Furthermore, the use similarity metrics which explicitly account for structural correspondences between simulated and reference data may also enhance the objective function.

**Marginal parameter distributions reveal optimal value-ranges** • Looking at the distribution of parameter values for the best samples tells us about “preferred” values for each parameter, for which the corresponding networks produce dynamical activity most similar to MEG resting-state data. Fig. 9 shows a comparison between the marginal parameter distributions computed independently for each of the four AC matrices. These distributions correspond to the 90^th^ percentile of all evaluated samples (ranked according to their similarity score). The narrower the distributions, the stronger the preference for a specific parameter value. And the more overlap between distributions, the better the consensus across experiments with different connectivities.

**Figure 9:**
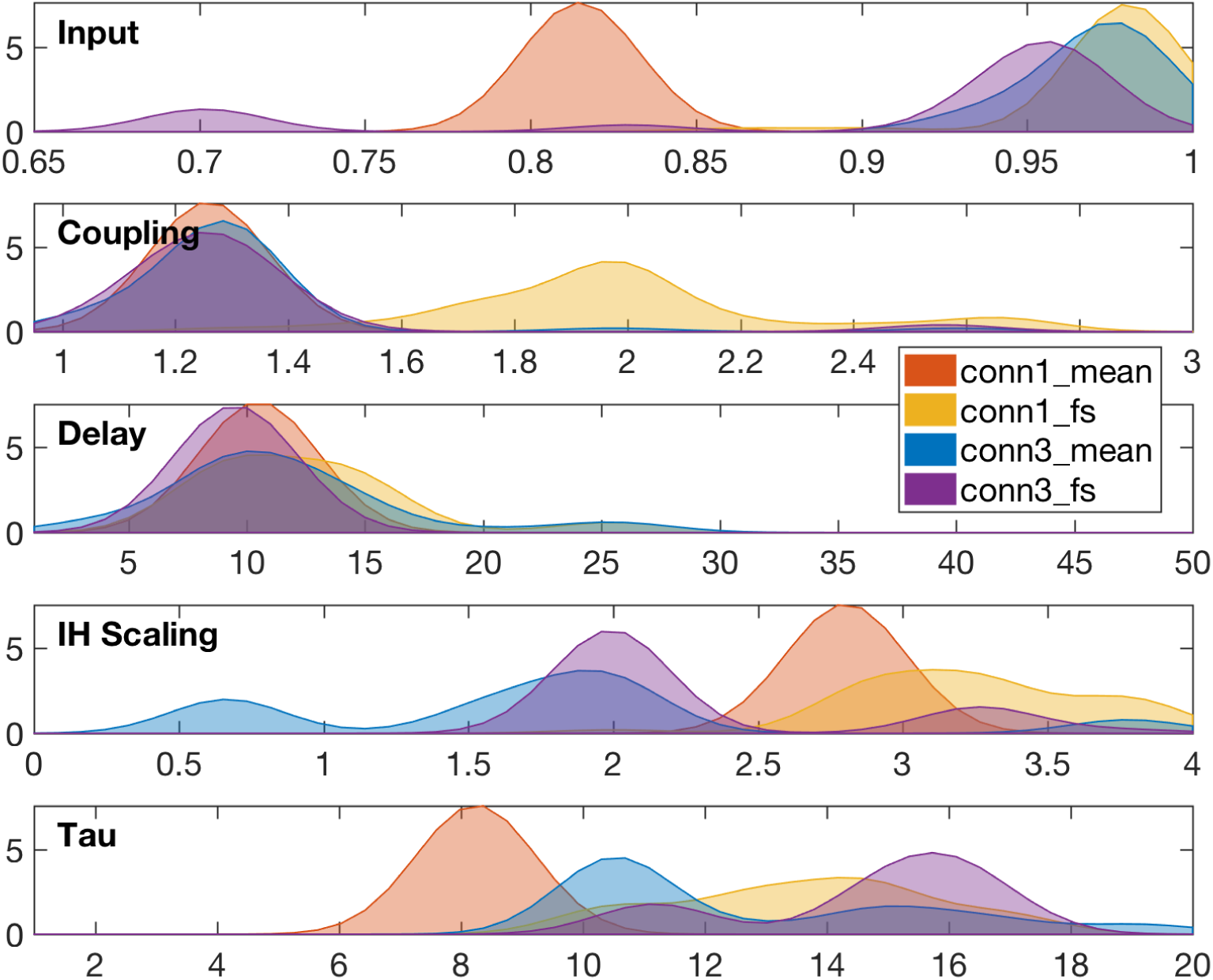
Marginal parameter distributions corresponding to the 90th percentile of all evaluated samples (*i.e.* using the objective function defined in §2.3.4), for each of the four AC matrices. Higher distribution values (y-axes) indicate ranges of parameters (x-axes) which were consistently associated with the best scores for a given AC matrix. **Input:** all but `conn1_mean` indicate that the excitatory input should be just below units’ oscillatory threshold. **Coupling:** all but `conn1_fs` indicate that coupling scale should be just above network oscillatory threshold. **Delay:** general consensus that average delay should be around 10ms. **Scaling:** no clear consensus, but all except `conn3_mean` indicate an upscale by a factor of 2 or more. **Tau:** `conn1_mean` centred around 8ms, and others above 10ms.

For example, we find a good consensus with regards to the first three parameters (input, coupling, delay), and in particular for the average network delay around 10ms, but the comparisons for the inter-hemispheric scaling *h* and characteristic time-constant *τ* are more mitigated. This is not surprising; the connectivity matrices control the interactions between the different brain regions, and structurally different networks should not be expected to agree on parameter values in general.

That being said, three out of the four AC matrices (all except `conn3_mean`) indicate clearly that the strength of inter-hemispheric connections should be increased at least two-fold. This is consistent with the known bias for shorter connections in probabilistic tractography, but it is also remarkable that we can estimate the amount of “missing” connectivity purely from simulations.

Finally, the results for the temporal parameters (average delay and time-constant) are somewhat surprising. We would not expect network delays to be lower on average than the characteristic time of variation within each brain region, because these delays are caused by axonal conduction over long distances, and local oscillations (caused by cycles of local excitation and inhibition) are not subject to propagation issues. This particular result might change with a more accurate estimation of the delays in our model (*e.g.* using tract-lengths from tractography instead of Euclidean distances), and may also be explained with further information about myelination information. Both of these avenues will be explored in future work.

**Conditional distributions reveal the local topography of the search space** • Here we take a deeper look at the best results obtained using `conn1_mean` connectivity. The optimal parameters correspond to a single point in the search space; to get an idea of the topography of the objective function around the optimum, we computed the conditional distributions of the GP surrogate on orthogonal slices going through that point. These slices are shown in Fig. 11.

A local maximum can be seen in the conditional surrogate coupling vs input (row 2 column 1), which indicates that the objective function is not unimodal. Note that this is by no means a complete picture; for example, it is impossible to know about local optima located elsewhere in the search space based on this information only. Instead, the partition tree from GPSO (not shown for brevity) can be used in combination with these conditional distribution, to identify local extrema and explore the topography of the search space around them.

Additionally, the marginally weighted means and standard-deviations of the similarity scores obtained during optimisation are shown on the diagonal of Fig. 11, computed within each dimension across *all* samples in eleven bins covering the corresponding parameter range. These statistics are consistent with the parameter distributions previously shown in Fig. 9, although we previously only considered the 90^th^ percentile of all samples.

**Region-wise correlations reveal poor correspondence in the frontal lobe** • The correspondence between the simulated and reference FC matrices shown in Fig. 8 can be explored further, by correlating each row of these matrices independently, in order to get a region-wise similarity score in each frequency-band. This comparison is illustrated in Fig. 10, by associating these correlations with a colour in each brain region and in each band. We find a very good correspondence across frequency bands in the temporal and occipital lobes, and systematically lower correlations in the frontal lobe, especially in the orbito-frontal cortex (OFC).

**Figure 10:**
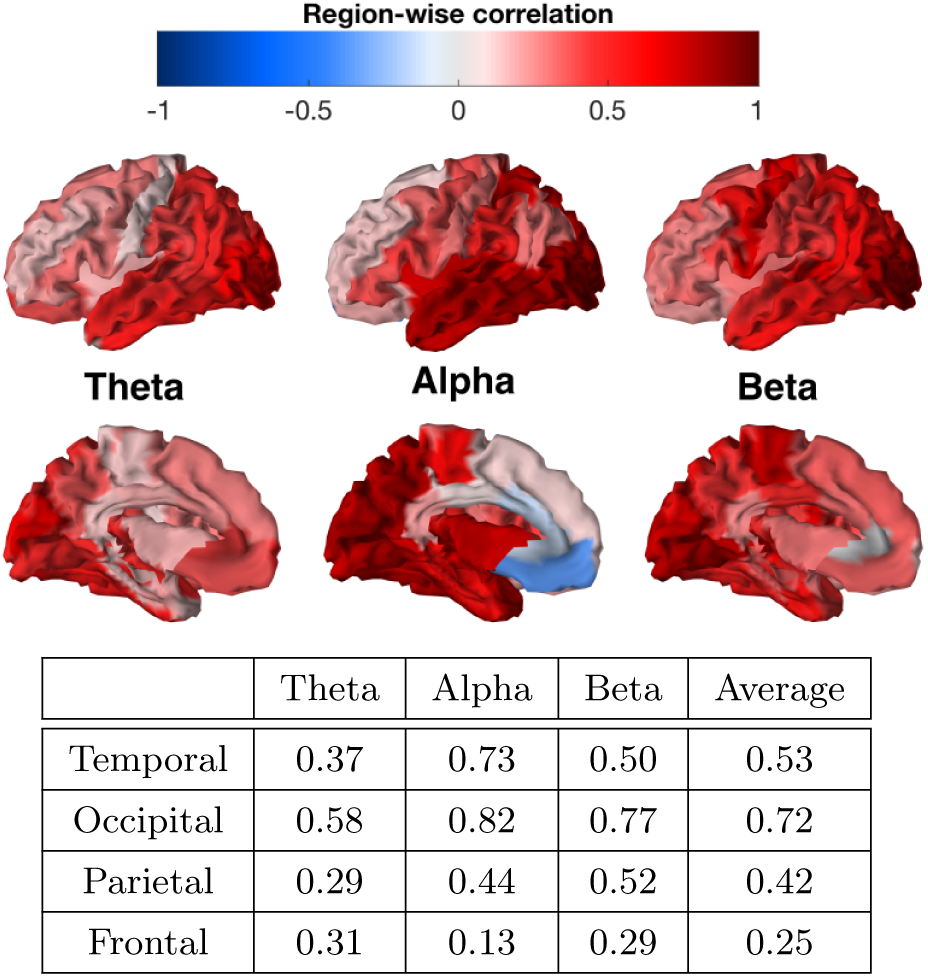
Region-wise correlation in each band, calculated between matching rows of simulated and reference FC matrices, for the best results obtained with `conn1_mean` connectivity. The average correlations within each lobe, for each band, are reported in the table below the surface illustrations. The correspondence between simulated and reference data is: very good in the occipital lobe; good in the temporal lobe, although driven mostly by the alpha band (>1.5 times better than other bands); consistently worse in the frontal lobe; and the average correspondence in the frontal+parietal lobes is twice as low as in the temporal+occipital lobes.

The signal-to-noise ratio in the OFC is known to be rather poor in MEG [23], but the fact that the bad correspondence extends throughout the entire frontal lobe may relate to the work of [10], which introduced gradients of excitatory inputs in the frontal areas, in order to account for higher dendritic spine counts compared with primary sensory areas. Such lobe-specific treatment can be easily introduced in our model (similarly to the inter-hemispheric scaling) and will be explored in future work.

Whether gradients of excitatory inputs improve the correspondence with real data or not, however, it is remarkable to be able to point to such specific modelling aspects, with reasonable confidence that no other configuration of the current system could yield a better result by tweaking the five parameters considered. These results tell us that a change to the *model* is required, and specifically one that will affect dynamics in the frontal areas. This type of information is invaluable, and demonstrates how GPSO can be used to inform modelling choices incrementally.

### 3.4. Discussion

To our knowledge, no other work in the literature attempted the systematic optimisation of LSBMs with dozens of brain regions, in order to model fast-paced electrophysiological dynamics, and controlling five (or more) parameters. The computational and theoretical complexity of these models (due to their non-linearity, but also their size and the presence of delays), combined with the richness of electrophysiological data calling for detailed objective functions leveraging the high temporal resolution, and the task of exploring parameter spaces as the number of dimensions increases (a.k.a. the curse of dimensionality), make the optimisation of LSBMs a truly difficult problem.

Our approach is different from the DCM method for network discovery [21], where the emphasis is put on inferring the *presence or absence* of structural connections, typically from fMRI data. For a given number of brain regions, this method considers all possible networks connecting these regions (that is, all possible combinations of edges), and proceeds to finding the network that is best supported by the observed data, as measured by the Bayesian model evidence, using generalised filtering [20]. Crucially, because it is impractical to list all possible networks beyond a handful of brain regions, let alone evaluate them, this method is made computationally efficient by exploiting the idea that it is sufficient to invert the fully-connected model in order to estimate the model evidence of any subnetwork. Furthermore, the method assumes that the posterior distribution over the connection strengths is multivariate Gaussian (the Laplace assumption); as such, it cannot represent accurately complex cost functions (*e.g.* with multiple modes, see Fig. 11), and in particular, only considers a single extremum during optimisation, which makes it prone to converging towards local extrema depending on initialisation.

**Figure 11:**
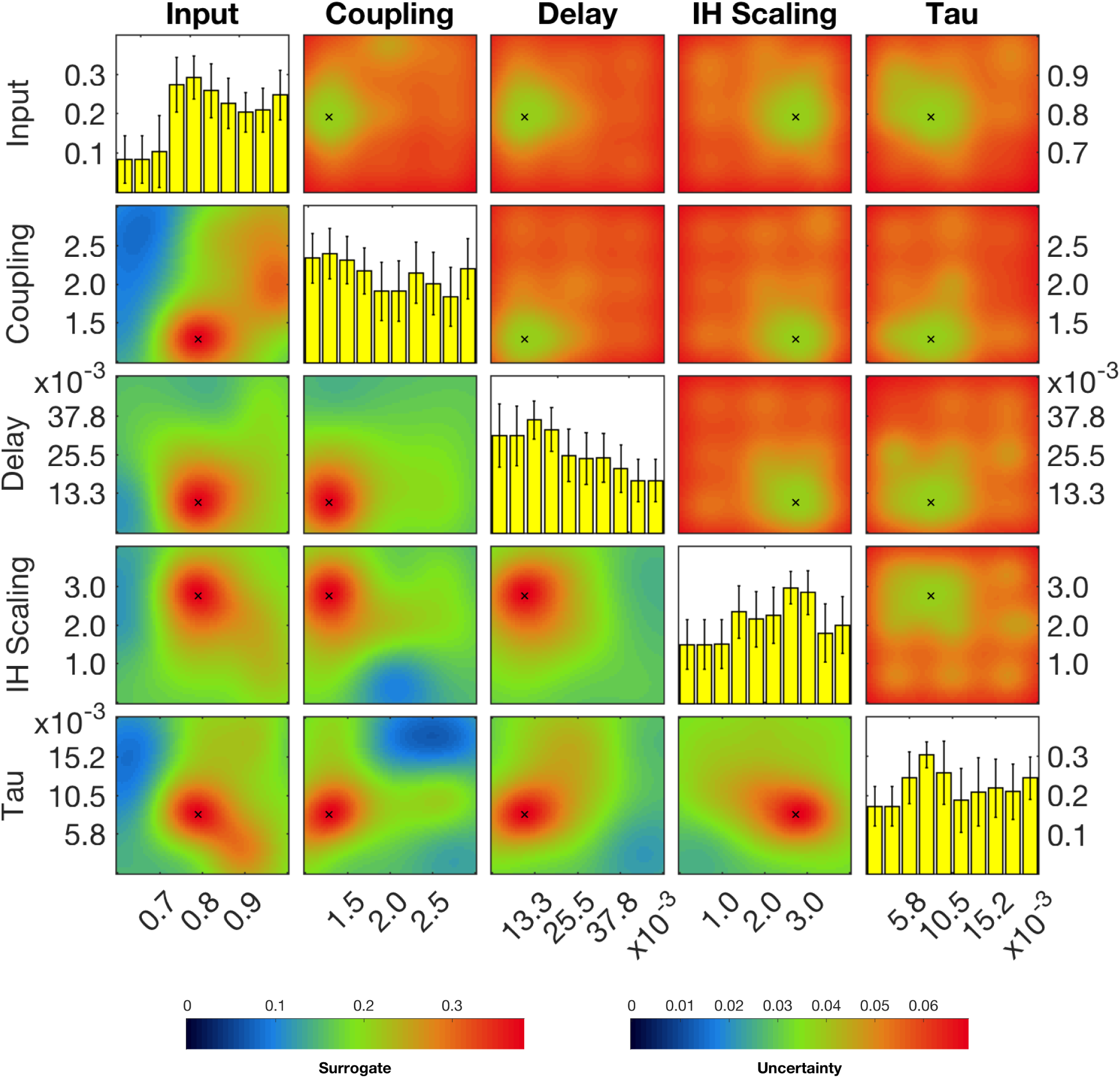
Conditional surrogate distributions (off-diagonal) and marginally weighted means and st-dev. (ondiagonal) around the best sample (black cross). These results correspond to the best experiment, using AC matrix `conn1_mean` (see Fig. 7). Note that y-axes at the top-left and bottom-right indicate similarity scores, whereas all other axes indicate parameter values. **Lower-triangle:** surrogate similarity (predicted mean) computed on orthogonal slices of the search space, going through the best sample for each pair of dimensions. **Upper-triangle:** associated surrogate uncertainty (predicted st-dev.) showing lowest uncertainty around the best sample, which is a good indicator of convergence. **Diagonal:** weighted mean and st-dev. of evaluated scores, calculated within each dimension across all samples. Higher bars indicate “preferred” values for the corresponding parameters (similar to the distributions shown in Fig. 9, but considering all samples).

In our case, the network is taken as the AC matrix estimated from diffusion tractography, and the emphasis is put on the Bayesian optimisation method proposed, which can be used to infer model parameters (up to a dozen in practice) with arbitrary objective functions encoding the dynamical features of interest. This method is capable of handling the computational burden associated with LSBM simulations in practice, and the presence of local extrema in the objective function. It does so by building a smooth *surrogate* of the objective function using a Gaussian Process, which is refined as the optimisation progresses, and exploited in order to prioritise the exploration of areas in the parameter space that are either unknown, or promising given the available evidence.

Nevertheless, there are a number of limitations currently associated with this method. **First,** it is not currently possible to systematically evaluate the convergence of the algorithm. This is mainly because at every iteration, multiple areas of the search space are being explored at multiple scales, which means that a lack of improvement in the best score obtained (typically a criterion for convergence) over several iterations is no guarantee that there will not be a substantial improvement at the next iteration. However, one can define several relevant termination criteria, such as: the number of evaluations of the objective function (our case), the number of iterations, the depth of the partition tree, etc. **Second,** it is worth noting that because we only ever select those nodes with maximal UCB in the partition tree (see Fig. 1), areas of the search space with lower expected scores are the last to be evaluated at each level of the tree, and therefore the resolution of the surrogate is lower there. This is an intended consequence of prioritising exploration in places of high expected reward, but it also means that the surrogate will in general not be reliable when the objective function is low; such is the price to pay for efficiency, this is not primarily an exploration method. **Third,** it is currently not possible to define priors over the parameter ranges in order to initially bias the search towards regions of known interest. Note that this cannot be done via the mean function of the GP, because hyperparameters are revised at each iteration, and that making the prior insensitive to hyperparameters would also make it insensitive to evidence accumulated by simulations, effectively corrupting the objective function as a result. It could however be done by introducing a third *type* of point (currently either evaluated, or GP-based, see §2.1.3), which would not be updated following hyperparameter updates, but would need to be evaluated before proceeding to exploration in an arbitrary small neighbourhood. This would essentially be equivalent to introducing “ghost nodes” arbitrarily deep into the partition tree, waiting to be discovered by subdivision. **Finally,** although this is purely a technical limitation, it is worth mentioning that the GP library we used (GPML [32]) is currently limited in the number of samples it can handle for regression; in practice, the regression becomes prohibitively slow beyond a few thousand samples, which means that we cannot reasonably explore parameter spaces beyond 10 dimensions. This can be solved indirectly, by selecting only a limited number of evaluated samples for training the GP; for instance, up to a certain depth in the partition tree, and randomly beyond that depth, up to a certain amount.

The two best results in our experiments, using `conn1_mean` and `conn3_fs` connectivity, indicate that inter-hemispheric scaling should be between two and three times as strong (see Fig. 9). Although these estimates should not be taken for granted without further validation (*e.g.* with different oscillatory models, or using fMRI reference activity), we want to highlight that they were obtained by optimising *structure* (the AC matrix) from *function* (band-specific FC); this is an exciting perspective offered by the method presented, with a different emphasis to previous work relating structure and function through biophysical models [38, 15].

To further elaborate on the validation of the results presented in §3.3: our experience suggests that small changes to the objective function can alter the results significantly (see Fig. 7); that different oscillatory models can lead to qualitatively different search-spaces (not shown); and the introduction of additional parameters can enable qualitatively different dynamics of the model. Furthermore, the frequency contents of the simulations (not included here into the objective function, but an important aspect of resting-state activity nonetheless) are affected by the heterogeneity of unit parameters across the network [10], and also most likely by the estimation of delays in the system; for instance, using tract-lengths instead of Euclidean distances, or including information about myelination.

Overall, the complexity of these systems makes it difficult to affirm with confidence that a given LSBM *cannot* produce dynamical activity with certain desired properties. However we argue that, for a given set of parameters, two models *can* be compared in terms of their performance with respect to an objective function (which encodes the desired dynamical behaviour) after optimisation. Provided that global convergence is achieved for both (we recall that the method proposed can handle local extrema), the comparison will still depend on the objective specified and on the chosen set of parameters, but will be valid under these conditions.

Previous related work in the literature such as [13, 9] did not formally employ optimisation methods to explore the capabilities of LSBMs, although they did study the effect of the connectivity strength and the average delay between brain regions (repectively *γ* and λ͐ in our model) on the simulated dynamics, by exhaustive grid search. We demonstrated that the method proposed can not only help speed-up this process considerably (see Fig. 6), but also allows to work in higher dimensional spaces by considering more parameters. This should enable more ambitious studies looking at the joint effects of structural and functional parameters on the simulated dynamics, and a principled comparison between different LSBMs in terms of measurable dynamical features (via the objective function), which will hopefully contribute to the ongoing development of a biophysical theory of brain activity.

## 4. Conclusion

We presented a Bayesian optimisation method capable of inferring the parameters of large-scale biophysical models (LSBMs) from imaging data. Using this method to optimise simultaneously five parameters, affecting both structural and functional aspects in delay-networks of 68 Wilson-Cowan oscillators, we were able to achieve the highest levels of expected correspondence with real resting-state MEG data across frequency bands, given the simulation time-lengths (see figures 8 and 5). Our results also suggest that inter-hemispheric anatomical connectivity, as estimated from diffusion tractography, may be underestimated by a factor 2 to 3, depending on the seeding and normalisation methods used. Furthermore, looking at region-wise correspondence in our best simulated results, we find systematically lower correlations in the frontal lobe, which indicates that further modelling work is required particularly in this area, perhaps in agreement with the work presented in [10]. Altogether, these results suggest that Gaussian-Process Surrogate Optimisation (GPSO) is an efficient and effective method for exploring the capabilities of LSBMs. It enables the exploration of high-dimensional parameter spaces (compared with the current state-of-the-art), which offers unprecedented insights into the relationship between structure and function in biophysical models of brain activity.

## 5. Acknowledgements

We would like to thank Matthew Brookes and his group for providing us with the MEG resting-state dataset of 28 healthy subjects. This data was collected as part of the Multi-modal Imaging Study in Psychosis (MISP) funded by grant MR/J01186X/1, and collected by Sian Robson and Emma Hall.

The Wellcome Centre for Integrative Neuroimaging is supported by core funding from the Wellcome Trust (203139/Z/16/Z).

J.H. is funded by the Oxford-Radcliffe DTC EPSRC Graduate Scholarship at University College, Oxford (EP/F500394/1). S.N.S. was supported in part by EPSRC (EP/L023067/1). M.W.W.’s research is supported by the NIHR Oxford Health Biomedical Research Centre, by the Wellcome Trust (106183/Z/14/Z), and the MRC UK MEG Partnership Grant (MR/K005464/1). S.J. is supported by the MRC UK (MR/L009013/1).

## Footnotes

1 For a GP with Gaussian likelihood kernel, the upper bound of a *p*% confidence interval on the expected value corresponds to *ς* = erfc^−1^(*p*/100), where erfc is the complementary Gauss error function.

2 Such delays are caused mainly by axonal conduction and synaptic transmission, both highly dependent on temperature, and range from hundreds of micro-seconds to tens of milliseconds at long-range [34].

3 The terms can be found at https://www.gnu.org/licenses/agpl-3.0. Briefly, any use of the code is permitted, without warranty, provided that copyrights are retained, and that any modification is made freely available under the same terms.

4 This link will be inactive until acceptance.

5 We know it is a fixed-point because we only choose inputs below oscillatory threshold (see Tab. 2).

